# Protective non-neutralizing mAbs Ab94 and Ab81 retain high-affinity and potent Fc-mediated function against SARS-CoV-2 variants from Omicron to XBB1.5

**DOI:** 10.1101/2023.09.29.560084

**Authors:** Arman Izadi, Magdalena Godzwon, Mats Ohlin, Pontus Nordenfelt

## Abstract

Antibodies play a central role in the immune defense against SARS-CoV-2. There is substantial evidence supporting that Fc-mediated effector functions of anti-spike antibodies contribute to anti-SARS-Cov-2 immunity. We have previously shown that two non-neutralizing but opsonic mAbs, Ab81 and Ab94, are protective against lethal Wuhan SARS-CoV-2 infection in mice. The protective effect was comparable to a potent neutralizing antibody, Ab59. Here, we hypothesized that, unlike the neutralizing antibodies, non-neutralizing opsonic antibodies would have a higher likelihood of retaining their function to the mutated variants, potentially functioning as broadly protective mAbs. Most of the mutations on the SARS-CoV-2 variants cluster on neutralizing epitopes, leaving other epitopes unaltered. We observed that neutralizing antibodies lost binding to Omicron. In contrast, seven non-neutralizing opsonic antibodies retained nanomolar affinity towards Omicron, BA.2, BA.4, and BA.5. Focusing on the two protective non-neutralizing antibodies Ab81 and Ab94, we showed that they maintain their strong reactivity even to XBB, XBB1.5, and BQ1.1. In the case of Ab94, interestingly, it even has increased affinity towards all variants except for XBB, which is comparable to WT. Finally, we show that Ab94 and Ab81 have potent Fc-mediated functions in vitro against the XBB and BQ1.1 and that combining the mAbs in a cocktail further enhances the effect. These results show that protective non-neutralizing mAbs such as Ab94 and Ab81 can be a viable strategy for anti-SARS-CoV-2 mAb therapies against current and possibly future SARS-CoV-2 variants and that opsonic epitopes could have implications for vaccine design.

## Introduction

SARS-CoV-2 has caused millions of deaths worldwide since it first emerged in 2019 ^1^. Monoclonal antibodies (mAbs) were successfully used as therapeutics early in the pandemic, ^2^. These antibodies disrupted the ACE2 receptor-spike receptor-binding domain (RBD) interaction, thus neutralizing the virus and protecting the host ^3^. However, with the emergence of variants, in particular Omicron, the antibodies mostly lost the ability to neutralize the virus ^4-5^. None of the clinically approved antibodies are functionally active against the emerging omicron subvariants XBB and BQ1.1 ^6^. The reason for this loss of function in these neutralizing antibodies (nAbs) was primarily the extensive mutations in the RBD present across the Omicron subvariants, at sites that most nAbs target. The emergence of these variants has led researchers to focus on generating pan-neutralizing antibodies ^7-8^. It remains to be seen whether this strategy will be successful against future variants of SARS-CoV-2.

Antibodies with potent Fc-mediated function have not been given the same level of attention as neutralizing antibodies. Recently, we have shown that two potent opsonizing antibodies, Ab81 and Ab94, targeting the RBD and N-terminal domain (NTD), respectively, strongly protect against the Wuhan strain in transgenic K-18 mice ^9-10^. Notably, these antibodies are non-neutralizing (nnAbs), mediating their protective effect through Fc-mediated effector functions. Other emerging findings with convalescent sera or neutralizing antibodies with modified Fc-effector functions show that Fc-effector functions are crucial for anti-SARS-CoV-2 viral control ^11-14^. The protection granted by vaccines against severe disease by mutated variants like Omicron can be partially explained by intact antibody Fc-effector functions directed against non-RBD sites ^15^. Thus, a viable strategy complementing the pan-universal neutralizing antibody approach could be to generate opsonic antibodies against conserved epitopes, avoiding the RBD- and NTD-sites targeted by most neutralizing antibodies.

SARS-CoV-2 variants have been heavily mutated. Bioinformatics analysis showed that the mutations occurring in the RBD and NTD cluster around neutralizing epitopes, leaving many other epitopes unaltered. This localized mutagenesis led to our hypothesis that opsonizing antibodies could be used for targeting these escape variants. To validate the approach of using opsonizing anti-spike antibodies as a potential therapy against Omicrons sub-variants and, in extension, future variants, we tested the reactivity of 8 clones generated against the Wuhan strain against several variants ^9^. We showed that of our 8 antibody clones, generated previously against the Wuhan strain, 6 retain unaltered binding against the Omicron, BA2, BA4, and BA5. Interestingly, only the neutralizing antibodies targeting the RBD lost binding. We further analyzed the affinity of our two protective clones 81 and 94 against the more recent variants XBB, BQ.1.1, and XBB.1.5, where we observed that they retain their nanomolar affinity to the spike protein. Finally, we show that both subclasses IgG1 and IgG3 of clones 81 and 94 show strong Fc-mediated phagocytosis against the newer XBB and BQ1.1 variants. A cocktail approach combining Ab81 and Ab94 generated the most potent functional output. The result shows a clear potential of using opsonizing non-neutralizing mAbs as a therapeutic strategy against the current and possibly future variants.

## Results

### SARS-CoV-2 variants primarily mutate in neutralizing epitopes of the NTD

In previous work, we established a set of monoclonal antibodies (mAbs) from B cells of previously SARS-CoV-2-infected subjects (**Fig. 1A**). We found that six mAbs (Ab11, 36, 66, 77, 81, and 94) were non-neutralizing but opsonic and capable of triggering phagocytosis. Two mAbs (Ab57 and Ab59) were both neutralizing and opsonic. The non-neutralizing Ab94 and Ab81 have both been shown to be protective against stringent infection (10^5^ PFU) of Wuhan WT strain in hACE2-K18 mouse model. In the case of Ab94, protection has been demonstrated both in a therapeutic and prophylactic model (**Fig. 1A**). However, whether Ab81 and Ab94, which bind to the RBD and NTD, respectively, would retain binding and protective function against the heavily mutated spike variants is unclear. It is well known that RBD mutations occur in neutralizing epitopes and sites relevant for ACE2-Receptor binding ^4-6^. On the NTD there exist five common sites which anti-NTD neutralizing antibodies target: residues 14-26, 67-79, 141-156, 177-186, and 246-260 (N1-N5 loops)^16-17^. Furthermore, residues 14-20, 140-158, and 245-264 have been identified as antigen supersites for anti-NTD nAbs ^16-17^. Epitopes outside the neutralizing residues are considered non-neutralizing and potentially opsonic.

**Figure 1.**
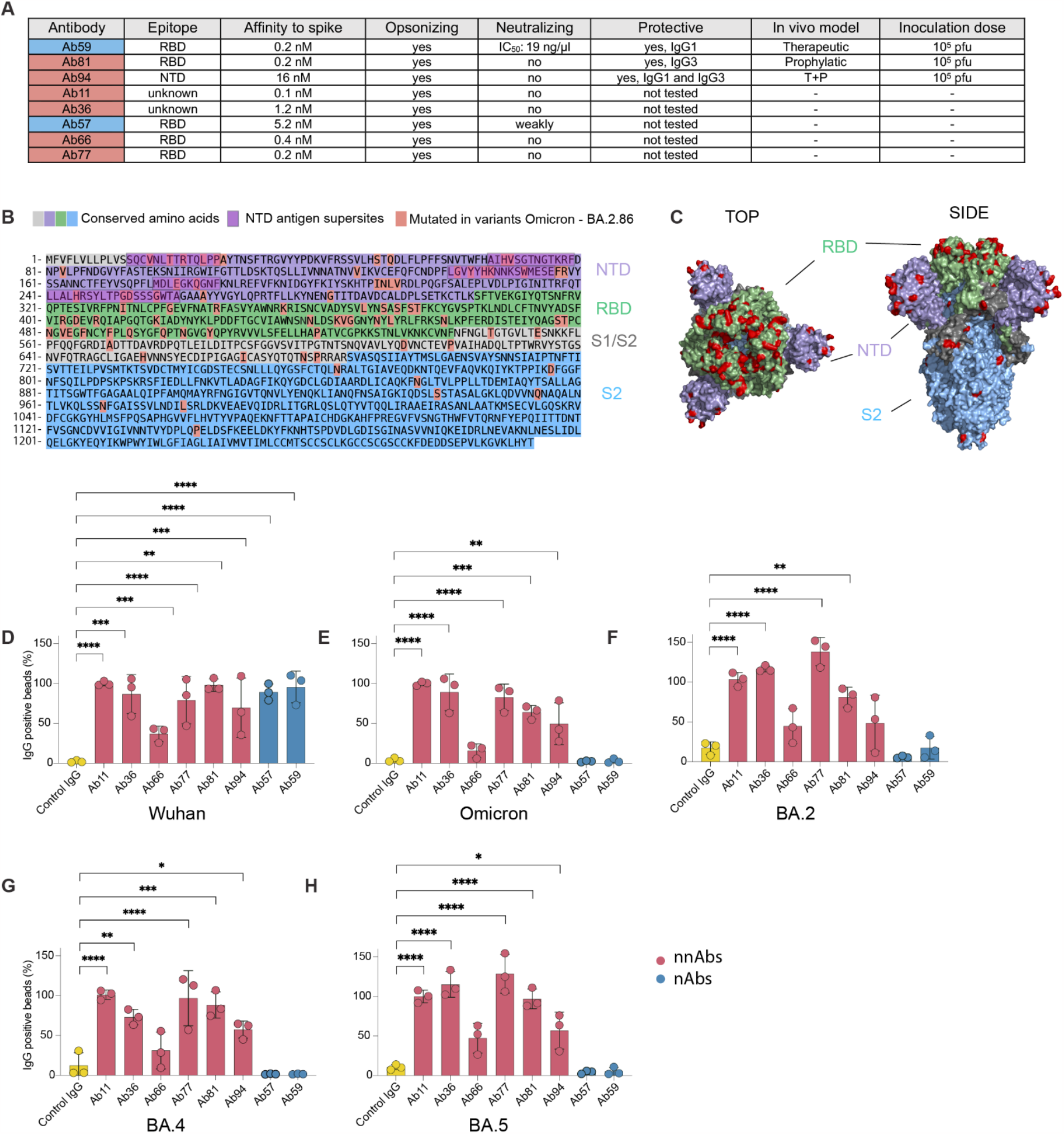
Protective non-neutralizing antibodies Ab81 and Ab94 retain binding against Omicron and sub-variants up until BQ.1.1, XBB, and XBB.1.5. **A** Table showing the characteristics of each antibody clone in terms of neutralizing ability, affinity to spike protein, opsonizing ability, and if tested and protective in an animal mode. **B** Spike amino acid sequence divided into the different domains (NTD, RBD, S1/2 and S2 domain). Mutations emerging from omicron to BA.2.86, according to the GSAID database are highlighted in red. **C** Spike trimer protein model with the RBD colored in green, NTD in purple, and the S2 domain in light blue. PDB ID: 6vxx^43^. Amino acid sequence of the spike protein of the Wuhan strain with mutated residues, seen also in B, highlighted in red on the spike model. **D** Wuhan spike protein was conjugated to microsphere beads and used as a model for SARS-CoV-2 virions. The spike-coated beads were opsonized with anti-spike mAbs. Detection of IgG-positive beads was done by adding an anti-IgG secondary, which was labeled with Alexa Fluor 488. The binding of anti-spike mAbs to spike-coated beads was done by flow cytometry. The Y-axis shows the percentage of IgG+ beads. The red-colored antibodies are non-neutralizing (nnAbs), and the blue are neutralizing antibodies (nAbs). Figure **D** depicts the reactivity of anti-spike antibodies against Wuhan, **E** against Omicron, **F** against BA.2, **G** against BA.4, and **H** against BA.5. In **D**, the percentage of binding was normalized to the background of Wuhan binding to Ab11. Statistical analysis was performed with one-way ANOVA with multiple comparisons against IgG control and corrected with Dunne’s correction test. ** denotes P-value < 0.01, * denotes P-value < 0.05 and P-value > 0.05 is ns.

We decided to look at all mutations in the relevant variants from Omicron up to EG.5.1 and BA.2.86. Interestingly, although the spike protein has undergone extensive mutations in both the RBD and NTD, most mutations occur in the neutralizing sites. In contrast, non-neutralizing sites generally have few mutations (**Fig. 1B-C, Supp. Fig. 1A-B**). The Omicron variant added mutations in the N2 loops (A67V, del69-70) but also in more distal sites in the NTD not associated with neutralizing mAbs (T95I, del211, L212I, and INS214EPE). The Omicron BA.2,4 and 5 subvariants added more mutations in the N1 (T19I, L24S, del25/27) and N3 loop (G142D). Only one mutation was added outside the neutralizing loops (V213G) (**Supp. Fig. 1C**). It is evident that most of the Omicron mutations occurred in the neutralizing epitopes in the N1-N3 loops and some mutations on residues 211-214, which are not associated with neutralizing antibodies. The SARS-Cov-2 variants further mutated in BQ1.1 and XBB, with a majority of the NTD mutations in the N2-N4 neutralizing epitopes (V83A, del144, H146R, Q183E) and one mutation between the N4-N5 loop (V213E) (**Supp. Fig. 1D**). We looked at the more heavily mutated variants XBB.1.5, 1.16, 1.19, 2.3x, CH1.1 and EG.5.1. It was even more clear that there is evolutionary pressure to mutate in the neutralizing epitopes with mutations primarily in the N3 (K147E, W152R, F157L), N4 (E180V) and the N5 loop (G252V, D253G, G257S) (**Supp. Fig. 1E**). EG.5.1 added a mutation between the N1-2 loop (Q52H). Finally, the newly emerged BA.2.86 has continued to add mutations in, or close to, the neutralizing epitopes: in the N1 (R21T), N3 (F157S, R158G) and N5 loop (H245N and A264D). While the other mutations from BA.2.86 correspond to non-neutralizing sites such as S50L, V127F, 211del, L212I, and L216F. Taken together, most of the mutations in the SARS-CoV-2 variants concerning the NTD end up in the neutralizing epitopes N1-N5 and some, with unknown significance, outside these loops, primarily in residues 211-216 (**Fig. 1B**). Thus, antibodies targeting residues outside of these loops can possibly retain their binding and function against these mutated variants. However, even if these mAbs bind to non-mutated epitopes, they might lose binding and function due to allosteric changes from other mutations. We decided then to move on to see if our protective Ab94 and Ab81 retain binding to their non-neutralizing but protective epitopes in the mutated SARS-CoV-2 variants.

### Protective non-neutralizing antibodies Ab81 and Ab94 retain binding against Omicron and sub-variants up until BQ.1.1, XBB, and XBB.1.5

Although we do not have complete information on the epitope sites for our anti-RBD nnAbs (clones 11, 36, 66, 77, and 81), we wanted to use these as well to test the hypothesis of using nnAbs to target, theoretically, less mutation-prone epitopes. In addition, we wanted to see if the nAbs Ab59 and Ab57 maintain their binding to the mutated variants. We used spike-coated microsphere beads to assess the reactivity of our mAbs against the Wuhan spike protein ^9-10^, by incubating the beads (coated with Wuhan, Omicron, BA.2, BA.4, and BA.5 spike protein) with our anti-spike antibodies. We assessed reactivity by adding fluorescent anti-Fab secondary antibodies (**Supp. Fig. 2A**). We analyzed IgG binding to the spike-coated beads with flow cytometry as done previously ^9-10^. All clones were analyzed as IgG3 except for Ab11, which was in the IgG1 subclass since it exhibited diminished affinity in the IgG3 subclass prior. Of the eight clones, which are strongly reactive to the Wuhan spike protein (**Fig. 1A, Fig.1D**), six showed strong binding to the Omicron variant (**Fig. 1E**). Interestingly, only the nAb clones (Ab57 and Ab59) were those that lost binding to the Omicron variant. Contrary to those, the nnAbs 11, 36, 66, 77, 81, and 94 retained strong reactivity. The same result was seen for the Omicron sub-variants BA.2, BA.4, and BA.5 (**Fig. 1E-H)**, reinforcing that opsonic antibodies can potentially bind to conserved epitopes while neutralizing antibodies are at significant risk of losing binding and function.

To further investigate the binding kinetics of our clones towards the spike variants, we performed surface plasmon resonance assays (SPR) with the multivalent spike trimer as done previously ^10^. For these assays, we chose to focus on clones 66, 77, 81, and 94 in the IgG3 subclass since these clones were shown to have potent *in vitro* Fc effector functions previously in this subclass ^10^. The results showed that the apparent dissociation constant (K_D_) and association rate constant (k_A_) are equivalent for all tested clones when comparing the reactivity towards the Wuhan spike protein and the variants Omicron, BA.2, BA.4 and BA.5 (**Fig. 2A-D, Fig. 2E**). Ab66 was an exception which showed reduced binding to BA.5 (3.75 nM) compared to WT (0.65 nM). More interestingly, Ab94 showed stronger binding to the mutated variants compared to WT (for instance, 2.5 nM to Omicron while 8.5 nM to WT), ranging from 3-fold better to more than 100-fold (BA.2) (**Fig. 2D-E**). Ab81 showed sub-nanomolar binding towards all variants tested. We further analyzed the avidity of clones 81 and 94 against the newer variants BQ.1.1 and XBB. We focused on these two clones since they were both shown to be protective against a lethal Wuhan infection in a K18-hACE2 mouse model ^9-10^. These SPR experiments showed that both clones retain a high affinity towards BQ1.1 and XBB (**Fig. 2F-G**). Ab81 shows some reduced binding towards BQ1.1 (1 nM) and sub-nanomolar to XBB (0.35 nM). Ab94 binds with similar avidity to XBB as to the Wuhan spike (k_A_ and K_D_) but, interestingly, binds 5-fold better to BQ1.1 following the similar pattern seen before with Omicron and its variants (**Fig. 2F-G**). Finally, we analyzed the binding ability of clone 81 against the RBD of XBB 1.5 due to its mutation in the RBD site F486P. This experiment showed a similar result that Ab81 has a sub-nanomolar affinity towards even this mutated variant of XBB (**Fig. 2F-G**) comparable to the affinity to Wuhan RBD measured previously (0.2 nM)^10^. We did not analyze Ab94 for this variant since the difference in mutational profile between it and XBB is identical when the NTD is concerned. In summary, the results show that the opsonic nnAbs retain strong binding to variants that emerged from Omicron while the nAbs clones did not. Finally, the protective clones Ab81 and Ab94 seemingly show high-affinity binding against these heavily mutated spike trimer variants. The highly protective Ab94 seems to bind even better to the mutated variants, except for XBB, which is comparable to the WT spike. This data reinforces the hypothesis that protective nnAbs, such as Ab81 and Ab94, can maintain strong reactivity towards mutated SARS-CoV-2 variants while nAbs lose their function.

**Figure 2.**
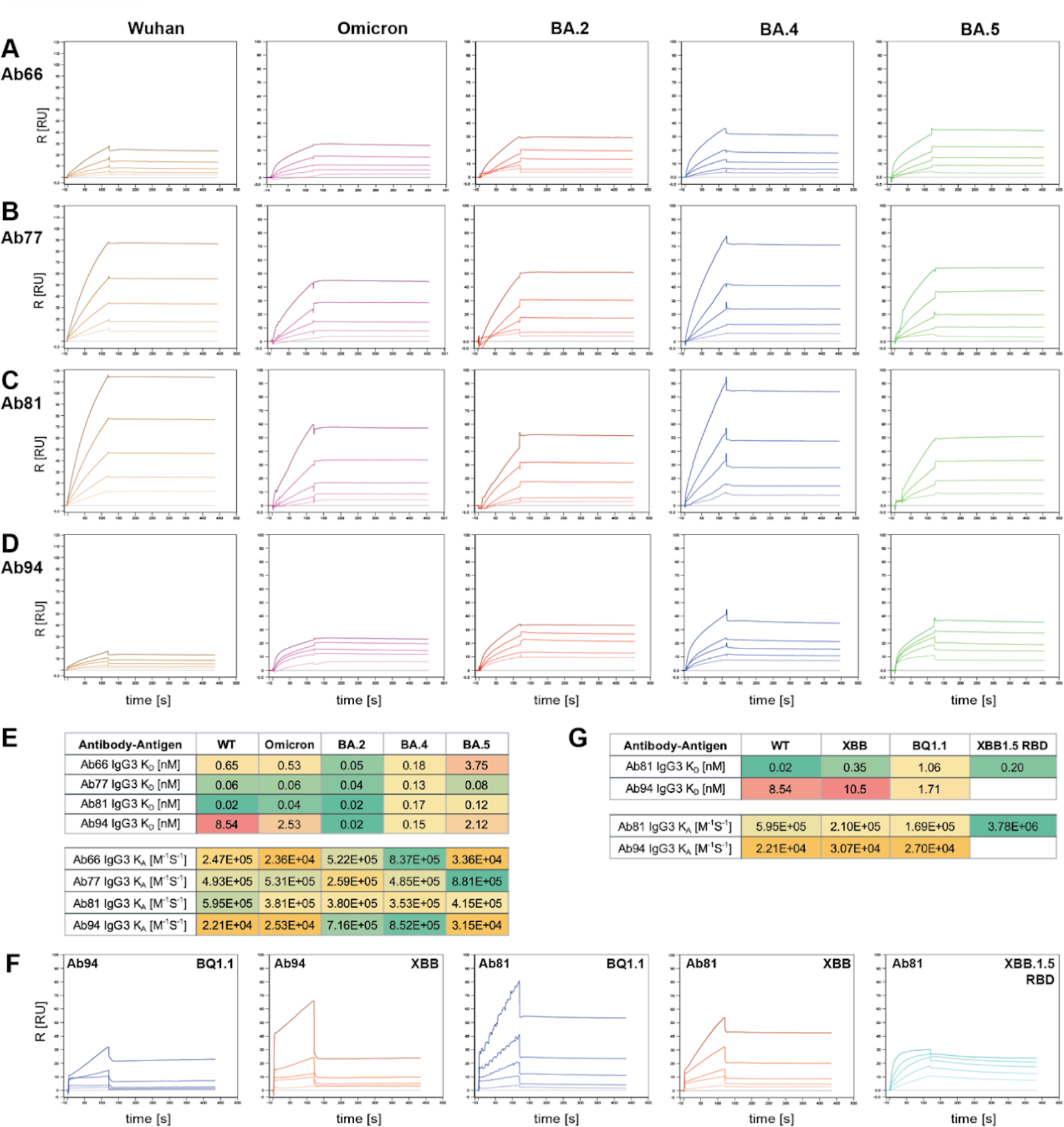
Binding kinetics with surface plasmon resonance reveal intact nnAb binding to spike variants. Surface plasmon resonance (SPR)-based binding kinetics assay with nnAbs against the spike trimer (WT, Omicron, BA.2, BA.4, and BA.5) as ligand. IgG concentration was kept constant on the sensor chip. In **A** IgG3 Ab66 is used in **B** IgG3 Ab77, in **C** IgG3 Ab81 and in **D** IgG3 Ab94. The spike concentration used ranged from 20-1.25 nM. **E** Table with K_D_-values and k_A_ for the nnAb antibodies against the spike variants. **F** SPR kinetic assay with clones 81 and 94 against XBB and BQ.1.1 spike trimer and XBB.1.5 RBD. **G** Table with K_D_-values and k_A_ for the nnAb antibodies against the spike variants.

### Both IgG1 and IgG3 subclasses of clones 81 and 94 show potent Fc-mediated phagocytosis against the BQ1.1 variant

Having established our clones’ binding properties, we decided to investigate if this translates to intact *in vitro* effector functions in the form of Fc-mediated phagocytosis. We opted to test their function on BQ.1.1 spike trimer-coated microsphere beads. We have previously shown that our mAbs are opsonic in a flow cytometry-based phagocytosis assay of spike beads with the THP-1 cell line ^9-10^ and that the results are relevant for biological protection *in vivo*. First, we tested the individual mAbs in both IgG1 and IgG3 subclasses for clones 81 and 94. We looked at the percentage of cells that were bead-positive and the median signal for beads in this population, which is a metric for bead quantity (**Supp. Fig. 3A**). Since the phagocyte-bead interaction is dependent on these metrics, we calculated a phagocytosis score based on the percentage of cells that are bead positive multiplied by the median fluorescence signal (bead signal) of that population, as described in other work ^18-19^.

Interestingly, both subclasses of IgG1 and IgG3 for both clones show potent Fc-mediated function compared to the negative control (Xolair, human IgG1 anti-IgE). This was observed in the percentage of cells that are interacting with the BQ1.1 spike-coated beads (bead^+^ cells), with all single IgGs showing comparable performance in this metric (ranging from 20-22% bead^+^ cells) compared to the negative control (9% bead^+^ cells, **Fig. 3A**). Furthermore, we analyzed the amount of beads being phagocytosed in cells associated with beads by quantifying the bead signal. In this analysis, IgG3 Ab81 stood out as the most potent opsonin (mean bead signal of 18200), followed by Ab94 IgG3 (15100, **Fig. 3B**). Ab81 and Ab94 IgG1 outperformed the negative control as well with mean bead signals of 13600 and 13000 respectively compared to the negative control (12000). Finally, we analyzed the phagocytosis score to acquire a comprehensive overview of the bead-phagocyte interaction. This analysis shows that Ab81 IgG3 enhanced bead-uptake by the THP1-cells more than 4-fold compared to the negative control (4300 vs 1000) while Ab94 IgG3 showed similar activity with a 3.5-fold enhancement (3500 vs 1030, **Fig. 3C**). No difference was seen between the IgG subclass controls (**Fig. 3D**). The IgG1 versions of clones 81 and 94 were 2.7-fold more potent than the negative control, trailing their IgG3 variants (2800 and 2700 respectively). Taken together, the results show that the opsonic clones 81 and 94 have strong Fc-mediated function against the BQ1.1 variant and that the IgG3 versions are, as the case with Wuhan, more potent opsonins.

**Figure 3.**
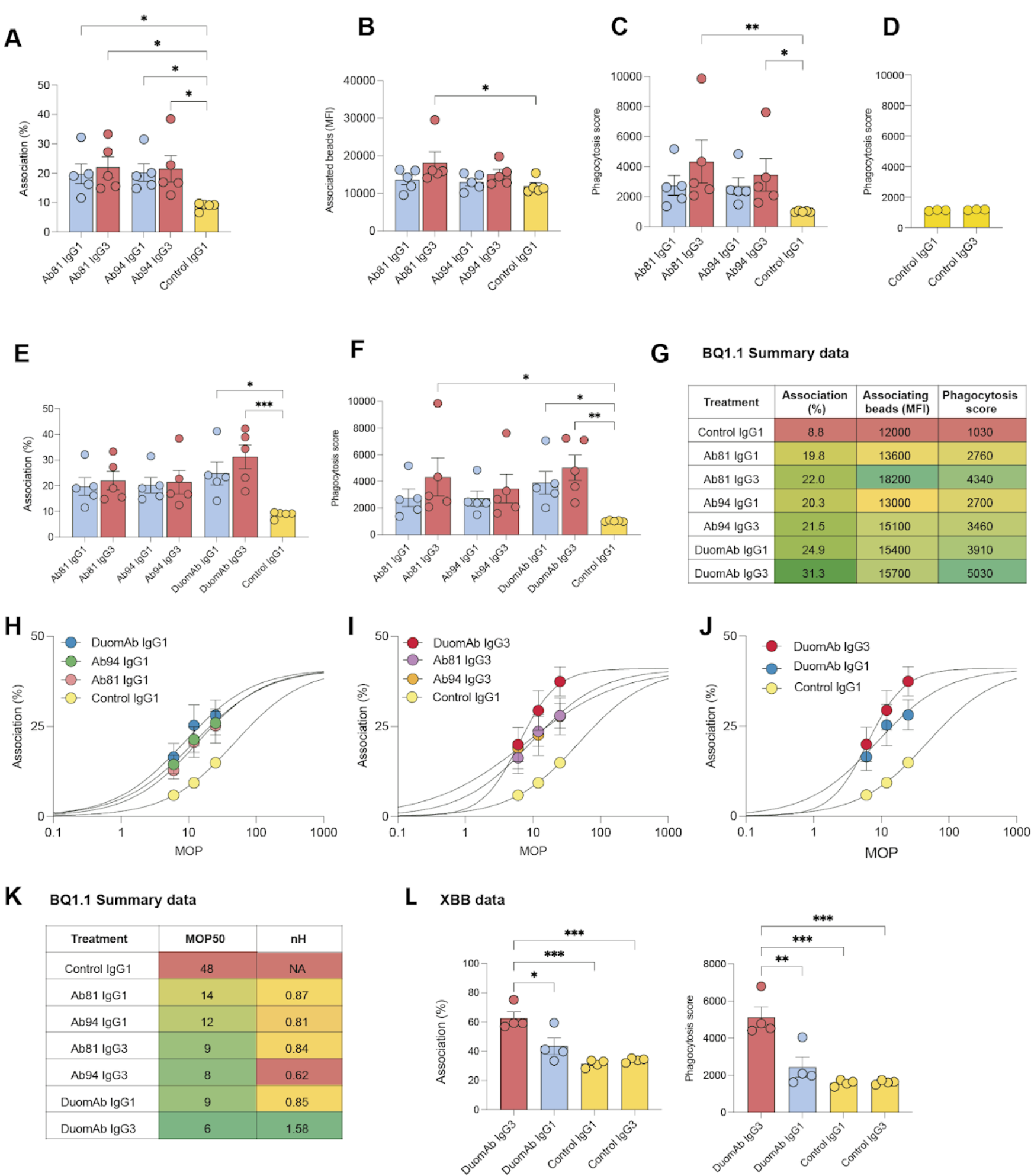
Both IgG1 and IgG3 subclasses of clones 81 and 94 show potent Fc-mediated phagocytosis against the BQ1.1 and XBB variants. Phagocytosis of spike-coated beads by the THP-1 cells was analyzed by flow cytometry. In **A-C, E-F** experiments were done at MOP 12.5 (100.000 THP1 cells, 1.25 million beads) with 2.5 ug/mL of antibodies to opsonize the spike beads. In short, THP1 cells were gated for bead signal (**Supp Fig. 3**) to generate % of bead positive cells (dubbed association) seen in A. The bead-positive cells were further analyzed for bead signal (Median fluorescence signal, MFI), a metric for the quantity of beads. By using association and MFI, a phagocytosis score was produced (% association multiplied by MFI/100). **A**. The percentage of bead-positive cells on the Y-axis with the treatments on the X-axis. **B** The bead signal (APC) for the bead-positive population on the Y-axis. **C** Phagocytosis score is shown on the Y-axis. **D** Shows the phagocytosis score for IgG subclass controls at MOP 25; no difference was observed. **E**. E Shows the percentage of bead-positive cells on the Y-axis at MOP 12.5 (100,000 cells, 2.5 million beads). **F** Depicts the phagocytosis score on the Y-axis at MOP 25. **G** Table summarizing the mean values from the experiments seen in A-F. **H-J** The graphs show association (bead^+^ cells) as a function of MOP (ranging from 25-6.25), and an EC_50_ was generated by a non-linear regression analysis (named MOP_50_). **H** Shows the differences between IgG3 single mAbs vs. cocktail. **I** Comparison of the differences between IgG1 single mAbs vs. IgG1 cocktails. **J** Shows the differences between IgG1 and IgG3 cocktails. In **K**, the MOP_50_ for each treatment is shown in addition to the Hill-slope value. In **L**, association and phagocytosis score by THP1 cells incubated with XBB spike-coated beads at MOP 25 is shown. In A-C, E-K, data are from 5 independent experiments. In D, it’s from three independent experiments, while for L, it’s from 4. In all figures, the mean is shown, and the error bars are SEM. Statistical analysis was performed with one-way ANOVA with multiple comparisons corrected with Dunne’s correction test for A-C and E-F compared against IgG control, while for L, comparisons were made for all treatments and corrected with Tukey’s correction test. ** denotes P-value < 0.01, * denotes P-value < 0.05 and P-value > 0.05 is ns.

### Cocktail of an anti-RBD and an anti-NTD mAb shows potent Fc-mediated phagocytosis in both IgG1 and IgG3 subclasses

In a previous study, we showed that combining a potent anti-NTD antibody (clone 94) with that of a potent anti-RBD (clone 81) generated a superior opsonic performance^10^. We named this cocktail DuomAb, and the IgG3 version of these mAbs showed the largest Fc-mediated effector functions against the Wuhan strain^10^. Having established potent function in both subclasses (IgG1 and IgG3) of Ab81 and Ab94 against BQ1.1 (the circulating variant in Sweden during the fall of 2022), we set out to test whether their potency could be enhanced by combining them in the same 2-antibody cocktail as done against the Wuhan strain (DuomAb) but against BQ1.1.

We performed similar experiments by incubating opsonized spike-costed beads with THP-1 cells. Only the DuomAb IgG3 outperformed the single mAbs at equivalent concentrations. DuomAb IgG1 and the individual mAbs Ab81 IgG1 and Ab94 IgG1 had very similar associating cells (24.9%, 19.8%, and 20.3% respectively, **Fig. 3E**). There was an increase in association with DuomAb IgG3 (DuomAb IgG3 = 31.3%, Ab81 IgG3 = 22.0%, Ab94 IgG3 = 21.5%, **Fig. 3E**). When looking at phagocytosis scores there was a difference for the DuomAb IgG1 vs. single IgG1 (DuomAb IgG1 = 3900, Ab81 IgG1 = 2800 and Ab94 IgG1 = 2700, **Fig. 3F**). Similarly as with percentage of bead^+^ cells, Duomab IgG3 had higher phagocytosis score compared to single IgG3 mAbs (DuomAb IgG3 = 5000, Ab81 IgG3 = 4300, Ab94 IgG3 = 3500, **Fig. 3F**). Compared to the negative control, Duomab IgG3 was the most potent opsonin, with a 5-fold higher phagocytosis score (**Fig. 3F-G**).

Furthermore, since the ratio of prey (spike-coated beads in this case) is a known factor influencing phagocytosis outcome ^20^, we decided to vary the ratio of prey to phagocytes, the multiplicity of prey (MOP), as described previously ^20^. We plotted the MOP against percentage associating cells and used non-linear regression analysis to generate the MOP_50_ value (equivalent to EC_50_) to compare the cocktails vs. single mAbs. A lower MOP_50_ value signifies an antibody that mediates more efficient phagocyte-prey interaction and is a more potent mediator of Fc-dependent phagocytosis. We observed that, while Ab81 and Ab94 IgG3 are both more efficient than the control (Ab81 IgG3 MOP_50_ = 9, Ab94 IgG3 =8, Control MOP_50_ = 48, **Fig. 3H**), they are both most effective when used in conjunction as Duomab IgG3 (MOP_50_= 6, **Fig. 3H**). Similarly as for the IgG3 variants, the IgG1 single mAbs outperformed the negative control (Ab81 IgG1 MOP_50_= 14, Ab94 IgG1= 12, Control MOP_50_ = 48, **Fig. 3I)** but were less efficient than their IgG3 counterparts. Here, the DuomAb IgG1 cocktail was more effective than both individual IgG1s (MOP_50_= 9). However, although Duomab IgG1 was more potent than its IgG1’s single mAbs, it was not more effective than Ab94 IgG3 and was inferior to both Duomab IgG3 and Ab81 IgG3, respectively **(Fig. 3H-J**). Further analysis indicates that DuomAb IgG3 exhibits a cooperative function with a Hill-slope constant (n_H_) above 1 (**Fig. 3K**). We did not observe such cooperative functions for the individual mAbs nor DuomAb IgG1. Finally, to extend our findings with the potent functions of DuomAb IgG3 compared to IgG1, we performed similar experiments with XBB spike trimer-coated beads. These results reinforced our findings with the Wuhan and BQ1.1 strain, with IgG3 DuomAb being highly opsonic compared to IgG1 DuomAb in terms of association (62.7% vs 43.5) and overall phagocytosis performance (5130 vs 2440) (**Fig. 3L**).

Taken together, our results show that both clones Ab81 and Ab94 exhibit potent Fc-mediated function against BQ1.1 and that engineering these clones in the IgG3 subclass and combining them in a 2-antibody cocktail generates even more potent functions against XBB and BQ.1.1. Since these nnAbs, especially Ab94, have demonstrated strong protectiveness against stringent infection in K18-hACE2 mice, our data suggests they could be promising candidates for therapy against BQ1.1, XBB and other variants.

## Discussion

Monoclonal antibody therapy against SARS-CoV-2 was successful in the initial phase of the pandemic ^2^. These antibody therapeutics neutralized antibodies that targeted specific RBD sites important for interaction with the ACE2-receptor. Unfortunately, mutated variants such as Omicron and its sub-lineages showed extensive mutations in these neutralizing epitopes, leading all clinically approved mAbs to lose binding and function^4-6^. Although many neutralizing mAbs generated initially against the Wuhan strain lost binding and function, researchers have successfully developed mAbs that neutralize the variants currently circulating ^7-8^. However, when testing these nAbs against emerging variants, several studies have shown a reduced neutralizing effect (IC_50_) compared to the Wuhan strain in the newer variants ^7-8, 21-22^. It appears likely that the trend will continue and that these nAbs will also eventually lose neutralizing capacity.

Neutralizing antibodies constitute a fraction of the adaptive antibody response generated against the SARS-CoV-2 virus. Most antibodies that bind to the spike antigen are not neutralizing ^23^. However, this does not necessarily imply that they do not serve any protective immune function. In previous work, we showed that all our antibody clones that bind to the spike protein promoted efficient phagocytosis through Fc-effector functions ^9^. The importance of Fc-mediated effector functions of anti-spike mAbs has for long not been the focus of the scientific community as an important immune mechanism against severe disease by SARS-CoV-2. For other viral pathogens such as adenovirus^24^, West Nile virus^25^, HIV-1^26^, RSV ^27-28^, Human CMV ^29,^ and Influenza virus ^30-32,^ antibody Fc-mediated effector functions are important for viral clearance. Thus, it is not surprising that more evidence is emerging showing that non-neutralizing antibodies mediate protective Fc-effector function against SARS-CoV-2. The evidence ranges from work with polyclonal antibodies from convalescent patients ^11,14^, modification of amino acids impacting Fc-effector function of neutralizing antibodies ^12^ to non-neutralizing antibodies ^6,10^.

Here, we show that two protective nnAbs, Ab81 and Ab94, retain high-affinity binding against variants up to XBB.1.5 and BQ1.1 and have strong Fc-mediated opsonic function against BQ1.1 and XBB. Although we do not know which exact epitope in the RBD clone 81 binds, or the NTD for Ab94, our experimental data shows that they have not been much affected by the current mutations in the RBD (up until BQ.1.1 and XBB.1.5). Both Ab81 and Ab94 were derived from Swedish patients in March 2020 ^9^, early during the pandemic. No variants of concern had yet surfaced ^33^. This means that these antibodies did not arise as a host response to any spike variants and that it is likely that the human immune system has been able to generate such protective nnAbs in response to SARS-CoV-2 infections all through the pandemic. This ability is likely common also in response to commonly used spike-based vaccines. The ability to generate protective antibodies against conserved non-neutralizing epitopes contributes to the ability of vaccination using early vaccine variants and prior, early infections to remain relatively effective in preventing serious disease, also following infection with emerging viral variants ^34^.

In our most recent work, Ab94 and Ab81 were both protective against an infection of authentic SARS-CoV-2 in K18-hACE2 mice in a prophylactic setting^10^. This is even more striking when considering that survival benefits were shown in a model suboptimal for human IgGs effector functions since human IgG (especially IgG3) promotes weaker phagocytosis by mouse phagocytes ^35-36^. Additionally, human IgG3 has a much shorter half-life in mouse circulation than human IgG1 ^37^. Therefore, the protective effects shown by these two mAbs could be underestimated in a human setting. Furthermore, Ab94 IgG1 was also shown to be protective in a therapeutic model, where it was comparable to clone 59, a potent nAb with IC_50_ of 19 ng/mL^9^. This comparison suggests that the protective benefit of nnAbs is comparable to that of nAbs. Notably, the mice were inoculated with a lethal dose of SARS-CoV-2 in both infection models, 10^5^ PFU ^9-10^. This dose is higher than that used in most studies that have shown the protective benefits of nAbs. Most researchers have tested the nAbs protective effect against a dose ranging from 10^3^ PFU to 10^4^ ^7-8,12,38-40,^ which are sublethal in comparison ^41^. Thus, not only do Ab81 and Ab94 retain binding and function against the new variants, but their Fc-mediated function has also been shown to be protective against death and weight loss in infection models where mice were challenged with a higher, and lethal viral dose compared to those commonly used to assess nAb protectiveness ^7-8, 2, 38-41^.

Although we recently showed that our opsonic mAbs are protective against the Wuhan strain in a K18-hACE2 mouse model, we did not perform similar experiments with the newer variants in this study. However, we did show that Ab94 and Ab81 maintain nanomolar affinity against the mutated variants and maintain their potent Fc-mediated effector functions. Taken together, our work with the nnAbs Ab81 and Ab94 shows the promise of using nnAbs as both prophylactic and therapeutic therapy against the current and future SARS-CoV-2 variants. Combining these nnAbs, or any other protective nnAbs, with nAbs could be one promising strategy to increase protective effects in vulnerable patients against the current and future variants. Although Ab94 and Ab81 are promising candidates, many conserved epitopes exist, especially in the S2 domain, which could yield further opsonic non-neutralizing antibodies. Finally, utilizing conserved epitopes capable of eliciting protective non-neutralizing antibodies would be useful for vaccine designs to combat future SARS-COV-2 variants and possibly other coronaviruses.

## Materials and Methods

### Cell culture and antibody production

Expi293F suspension cells were purchased from Gibco (ThermoFisher) and routinely cultured in 125 ml Erlenmeyer flasks (Nalgene) in 30 ml Expi293 medium (Gibco) in an Eppendorf s41i shaker incubator at 37°C, 8 % CO_2_, 120 rpm. Cells were routinely passed and split to a density of 0.5 × 10^6^ cells/ml every 3 to 4 days. The day before transfection, the cells were seeded at a density of 2 × 10^6^ cells/ml. The next day, cells were seeded at 7.5 × 10^7^ cells in 25.5 ml Expi293 medium. Transfection with heavy and light chain plasmids was performed using the Expifectamine293 kit (Gibco) according to the manufacturer’s instructions. The plasmids for the heavy chains were generated previously ^9-10^. Briefly, 20 μg of heavy and light chain plasmid, respectively, were mixed with 2.8 ml OptiMEM (Gibco) and 100 μl Expifectamine and incubated at room temperature for 15 minutes. Afterward, the transfection mix was added to the Expi293F cells. The following day, 1.5 ml of enhancer 1 and 0.15 ml of enhancer 2 (both from the Expifectamine293 kit) were added, and the cells were cultured for another 72 hours.

The cells were removed from the cell culture medium by centrifugation (400 x g, 5 min), and the supernatant was transferred to a new tube. To capture the IgGs from the medium, protein G sepharose 4 Fast Flow (Cytiva) was added to the medium and incubated end-over-end at room temperature for 2 hours. The beads were collected by running the medium bead mix through a gravity flow chromatography column (Biorad) and washed twice with 50 ml PBS. The antibodies were eluted using 5 ml HCl-glycine (0.1 M, pH 2.7). Tris (1 M, pH 8, 1 ml) was used to neutralize the pH. The buffer was exchanged to PBS using Amicon Ultra-15 centrifugal filters (Sigma) with a molecular cut-off of 30,000 Da. The concentration and quality of the purified antibodies were spectrophotometrically measured with the IgG setting of the Nanodrop (Denovix).

### Generation of Spike-coated beads

Wuhan Spike-protein was generated by transfecting Expi293F cells with 40 µg plasmid containing the gene for the Spike protein (CS/PP Spike encoding a secretable version of the protein was used to allow purification from cell culture supernatants), donated previously to us by Dr. Florian Krammer’s lab ^5^. Omicron (#158-40589-V08H26-100), BA.2 (#158-40589-V08H28-100), BA.4 (#158-40589-V08H32-100), BA.5 (#158-40589-V08H33-100), XBB (#158-40589-V08H40-100) and BQ1.1(158-40589-V08H41-100) was aquired from Nordic biosite. 25 µg of Spike protein was biotinylated according to the instructions of EZ-LinkTM Micro Sulfo-NHS-LCBiotinylation Kit (ThermoFisher). Then fluorescent (APC) streptavidin microsphere beads (63µl from stock) (1 µm, Bangs Laboratories) were conjugated with the biotinylated Spike protein (25 µg) according to the manufacturer’s instructions as done previously ^9-10^.

### Flow cytometry-based avidity measurements

The binding assays were performed in a 96-well plate, pre-coated with 200 µl of 2% BSA (in PBS) at 37 degrees for 30 minutes. 250,000 Spike-coated beads were used in all wells, with an antibody concentration of 1 µg per ml. The beads were opsonized at a volume of 100 µl in 1X PBS at 37 °C for 30 min on a shaking heat block. The wells were washed twice with PBS. To assess antibody binding to Spike beads, 50 µl of (1:500 diluted) a Fab-specific fluorescent secondary antibody (#109-546-097, Jackson ImmunoResearch) was used to create a fluorescent signal. The secondary antibody was left to incubate with the Spike-bead antibody complex at 37 °Cs for 30 minutes on a shaking heat block. 100 µl of PBS was added to the wells before analysis in the flow cytometer. The beads were analyzed using a Beckman Coulter Cytoflex flow cytometer, which acquired 10000 beads per sample. The data was processed using Flowjo. The gate for Spike beads was set based on forward and side scatter **(Supp. Fig 2A)**. The gate for Spike beads positive for antibodies was set based on reactivity to a non-reactive IgG control mAb.

### SPR kinetic assay

To study binding kinetics to Spike trimer or NTD we immobilized A High Capacity Amine Sensor chip (Bruker) was immobilized with Anti-human IgG (Fc) antibody (Cytiva BR-1008-39) at 25 µg/ml in 10 mM sodium acetate buffer pH 5 at flow rate 10 µl/min and contacting 300s. This was done in a MASS-16 biosensor instrument (Bruker). Running buffer PBS + 0,05%Tween20. Antigens were acquired from Nordic biosite (Omicron#158-40589-V08H26-100,BA.2#158-40589-V08H28-100,BA.4 #158-40589-V08H32-100,BA.5#158-40589-V08H33-100,XBB#158-40589-V08H40-100, XBB.1.5 #158-40592-v08h146-100, BQ.1.1 #158-40589-V08H41-100).The antibodies were diluted in PBS and injected over the surface for 90s at 10 µL/min. The running buffer was PBS with 0.01% Tween 20. All variants of the Spike trimer were analyzed at 20 nM to 1.25 nM concentrations. The XBB.1.5 RBD was analyzed at concentrations ranging between 20 nM to 1.25 nM as well. The antigens were injected at these concentrations and were allowed to interact with the sensor for 2 minutes, with flow rate 30 µl/min, followed by a dissociation for 6 minutes. After each cycle the surface was regenerated with 3M MgCl. All experiments were performed once. The data was analyzed using Sierra Analyser software version 3.4.3 (Bruker) program to determine apparent dissociation constants (K_D_) and constant rates (k_A_)

### Flow cytometry-based phagocytosis assays

THP-1 cells (Sigma-Aldrich) were cultured as described previously ^5^. In all experiments, 1×10^5^ THP-1 cells were used. In the experiments, the ratio of cells to beads ranged from 6.25 to 25 for the curves (MOP 6.25-25) and MOP 25 for the XBB experiments. For all experiments, the Spike beads were opsonized with 2.5 µg/ml of antibodies in a volume of 100 µl of Sodium media (pH adjusted to 7.3 with NaOH; 5.6 mM glucose, 10.8 mM, 127 mM NaCl, KCl, 2.4 mM KH_2_PO4, 1.6 mM MgSO4, 10 mM HEPES, 1.8 mM CaCl_2_). Opsonization was performed for 30 minutes at 37 °C on a shaking heat block (300 RPM) in a volume of 100 µl. During this incubation period, THP-1 cells were counted (using a Bürker chamber) and medium was exchanged from RPMI to Sodium medium. THP-1 cells (50 µl, 2×10^6^/ml) were added to each well and the cells were allowed to phagocytose beads for 30 minutes at 37 °C on a shaking heat block (300 RPM). The 96-well plate was then incubated on ice for 15 minutes and analyzed directly in a Beckman Coulter Cytoflex flow cytometer. A gate for THP-1 cells was set up based on their forward and side scatter. The gate for association was set with a negative control of cells only. The analysis stopped after 5000 events were captured in the THP-1-gate. The data was analyzed in the program Flowjo by setting similar gates as described above **(Supp Fig. 3A)**. The FlowJo-processed data was further analyzed in GraphPad Prism where the bead signal (APC-A) of the THP-1 gate association gate, percentage of associating cells and phagocytosis score were plotted to compare the different antibodies. Phagocytosis score was calculated based on the percentage of bead^+^ cells (associating cells) multiplied by the median fluorescence intensity (APC-A) of that population as described previously^18-19^.

### Statistical analysis

Statistical analysis was performed in GraphPad Prism. For comparisons between clones when more than two treatments were analyzed, a one-way ANOVA with multiple comparisons test was used, with correction for multiple comparisons with Dunnes’ test or Tukey’s, depending on the comparisons being made.

### Bioinformatics

Analysis of spike mutations was performed using the lineage comparison tool at GSAID described recently ^42^ by aligning the strains of interest with that of a reference strain B.1.

## Acknowledgments

We acknowledge funding from the Swedish Research Council, Crafoord Foundation, and Knut and Alice Wallenberg Foundations, and support from an infrastructure grant from the Faculty of Engineering, Lund University.

