## Supplemental Info for "Protective non-neutralizing mAbs Ab94 and Ab81 retain high-affinity and potent Fc-mediated function against SARS-CoV-2 variants from Omicron to XBB1.5"

**This PDF file includes:**

Figures S1 to S3

### Supplementary Figures 1-3

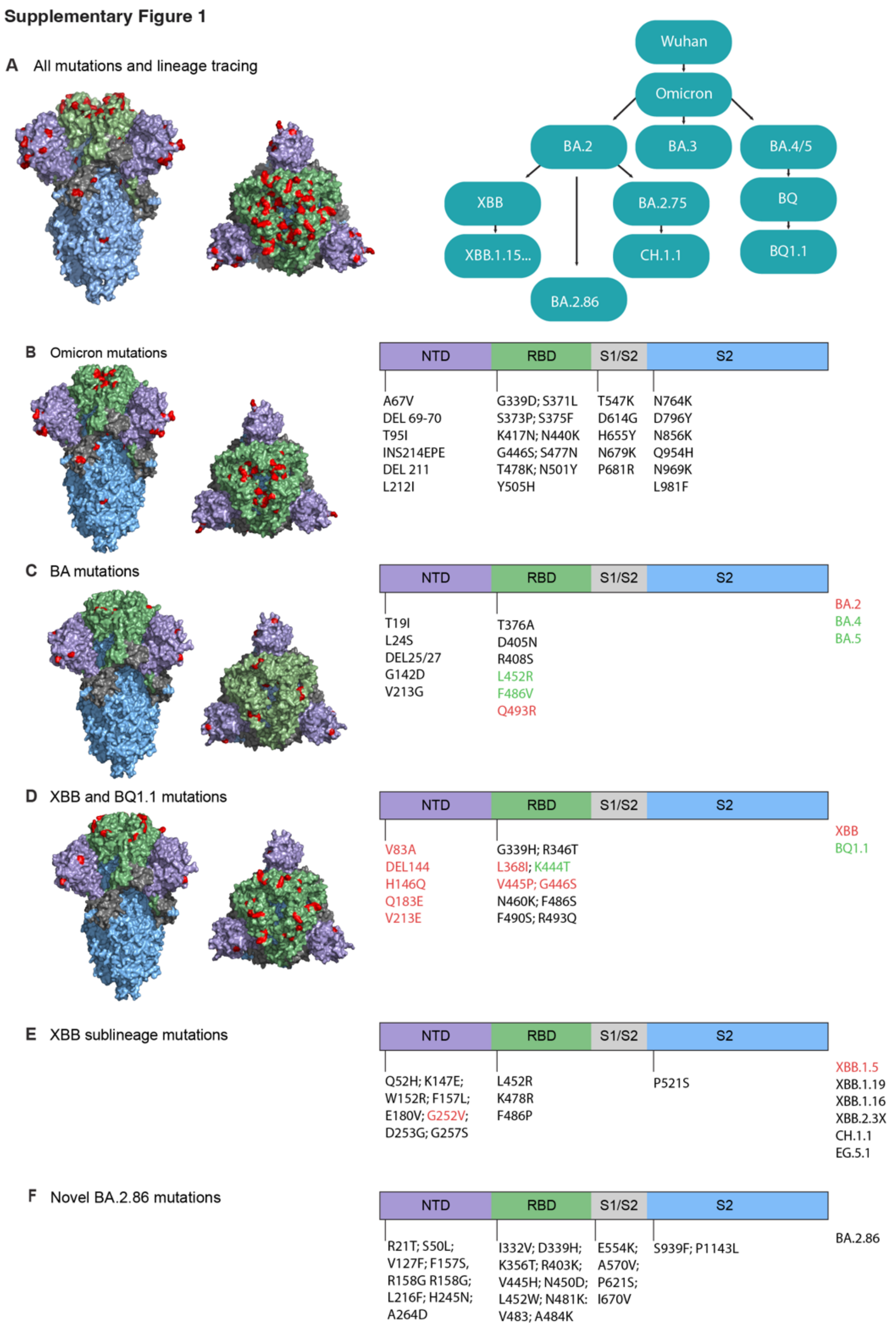


**Fig. S1. A** Visualization of all spike mutations on Wuhan Spike trimer. Lineage tracing of omicron subvariants is shown. **B-F** mutations in the respective domains of the spike protein by each variant. Spike trimer protein model generated with Pymol with the RBD colored in green, NTD in purple, and the S2 domain in light blue. PDB ID: 6vxx[1].

**Supplementary Figure 2**

A

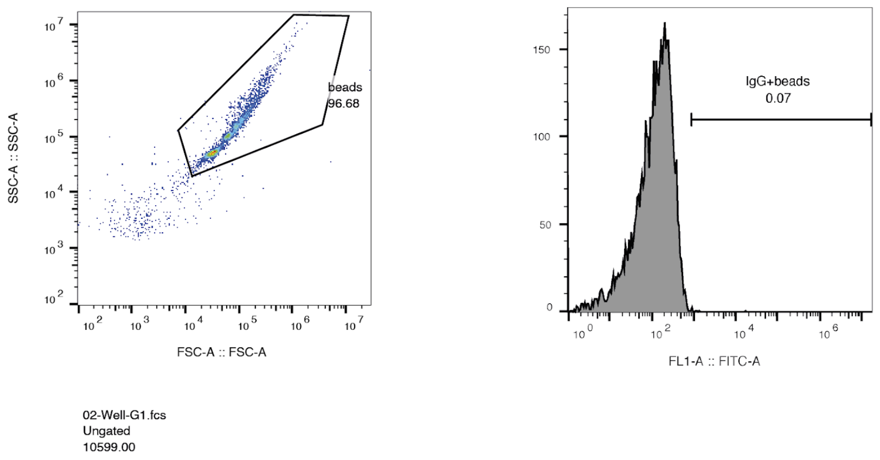


**Fig. S2. Flow-cytometry binding assay gating strategy.** A. Beads were gated for by size and granularity. Thereafter beads were gated for by IgG binding (FITC signal) using a negative control of beads only.

**Supplementary Figure 3**

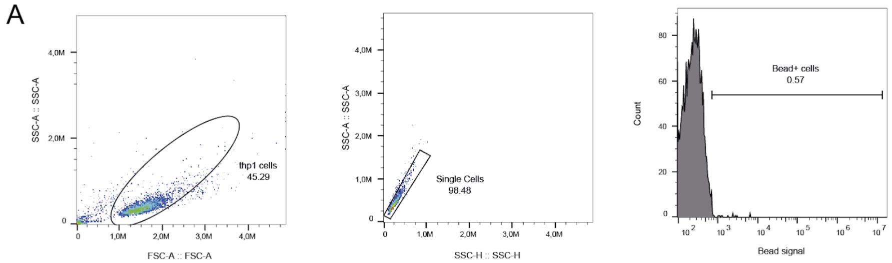


**Fig.S3. Phagocytosis assay gating strategies. A** THP1-cells were gated for by size and granularity, then a single cell gate was made based on granularity height and area to exclude duplicate events. A bead positive gate was drawn based on a negative control with cells only.
